# Learning from Machine Learning: advancing from static to dynamic facial function quantification

**DOI:** 10.1101/2024.03.28.584911

**Authors:** Sandhya Kalavacherla, Morgan Davis Mills, Jacqueline J. Greene

## Abstract

**Objectives:** We assess an open-source Python machine learning algorithm’s efficacy in image and video analysis of facial palsy (FP) patients.

**Methods:** Images and videos of 60 patients with varying FP severities performing standard movements were obtained from MEEI Facial Palsy database. Landmarks generated on images by the open-source algorithm (adapted from OpenCV and Dlib libraries) and Emotrics (standard for two-dimensional FP analysis) were compared. Considering the human eye as the standard for accuracy, three raters marked perceived errors in each algorithm’s tracking of five facial features. Cumulative error distributions between both algorithms were compared via normalized root mean square error. FP severity and facial feature-specific error rates were compared using ANOVA tests. Raters also analyzed open-source algorithm-generated video landmarks; similar statistical comparisons between open-source algorithms’ image and video-based analyses were performed.

**Results:** Cumulative error distribution between both algorithms’ image analyses was most robust for normal function; significant discrepancies were observed in mild/moderate flaccidity and nearly-normal/complete synkinesis. Both algorithms had similar error rates across all facial features (p=0.76) and FP severities (p=0.37). In the open-source algorithm’s video analysis, mild synkinesis (24.7%) and severe flaccidity (19.7%) had the highest error rates. Comparing image and video analyses generated by the open-source algorithm, video analyses had lower error rates across all FP severities (p<0.001).

**Conclusions:** We report for the first time the feasibility and relative accuracy of a Python open-source algorithm for dynamic facial landmark tracking in FP videos. The demonstrated superiority of landmark tracking with videos over images can improve objective FP quantification.

## Introduction

An objective, logistically feasible, automated and accurate method to quantify dynamic facial function has been a long-standing challenge in the care and management of facial palsy (FP) and facial nerve disorders.^1^ Clinician-based facial analysis scales have evolved from the House-Brackmann (HB) to the Sunnybrook and eFACE, but have limited objectivity^1–7^ with reported interobserver variability up to 64% for severe FP (HB III or worse).^8^ Patient-based surveys such as the FACE Instrument pose a similar limitation, although they also uniquely capture the individual’s experience.^7^ While artificial intelligence (AI) and machine learning (ML) hold promise for improved methods of automated quantification of facial function, most currently available ML algorithms are either limited to static image analysis (such as Emotrics or the auto-eFACE ^9,10^), require proprietary marketing software to estimate facial emotional expressions^11–15^ or were limited to a single surgical case report^16^ following surgical interventions. An objective, open-source, rigorous quantification of dynamic facial function in videos has remained elusive.^16–18^

Landmark errors represent a significant challenge for automated computer-aided facial analysis of either photos or videos in the FP population or those with neurologic deficits, as the ML algorithms were trained on large datasets comprised of subjects with normal facial function.^10,15–17,19–24^ Open-access software such as Emotrics^9^ based in Python ML-libraries (dlib, OpenCV) have made improvements to reduce landmark inaccuracies in static images of FP patients through allowing manual landmark adjustment of available datasets of FP patients and retraining ML algorithms on corrected images. However, employing manual adjustments to videos was deemed to be resource-intensive, error-prone and infeasible for dynamic video-based ML-facial analysis,^9^ especially given the small FP patient population and limited available data sets. However, open-source and freely available computer vision and ML algorithms enabling facial analysis of videos (and incorporating the same ML-libraries dlib, OpenCV) have been available through Python since 2015, albeit not fine-tuned for FP analysis.^20,25,26^ To our knowledge, the feasibility and accuracy of these open-source programs in video analysis of facial landmark recognition in FP patients has not yet been described. While inference of dynamic function from static images of a standard facial nerve exam is commonly reported^9,10,27^, videos can provide a more comprehensive picture of the nature of a patient’s facial function and thus serve as a rich resource to quantify dynamic facial function.

The goal of this study was to 1) compare the accuracy of landmark detection between Emotrics and Python open-source ML-algorithms in static facial image analysis, 2) to test the feasibility of utilizing the open-source Python ML-algorithm to analyze videos of FP patients and 3) to compare video landmark error rates with static image analysis. We provide the adapted Python code as a resource to promote further work in this area. This study lays the groundwork towards generating quantitative, objective and automated scores of dynamic facial function that can assess severity and track recovery, improving the clinical utility of AI-aided automated FP assessment.

## Methods

### Facial Palsy Database

We utilize the MEEI [Massachusetts Eye and Ear Infirmary] Facial Palsy Photo and Video Standard Set (https://www.sircharlesbell.com/), an open-source, prospectively gathered, standardized set of facial photographs and videos from 60 participants representing the entire spectrum from normal to flaccid and synkinetic (aberrantly regenerated) FP.^28^ Subjects enrolled in this standard set were categorized by their eFACE and HB score into the following: normal (96–100, HB 1), near-normal (91–95, HB II), mild (80–90, HB II-III), moderate (70–79, HB III), severe (60–69, HB IV-V), and complete flaccid (<60, HB VI) or nonflaccid FP (<60, HB IV). Complete demographic details are available in the original publication^28^; in brief, the average age of participants was 47±17 years, reported gender was 45% male, and Bell’s Palsy was the most common FP etiology. The images and videos were taken with a Nikon DSLR camera in studio lighting and against a standard blue background.

### Adaptation of Emotrics and Python ML-algorithms for Batch Processing

Emotrics is freely available as a downloadable GUI and with the source code available (https://github.com/dguari1/Emotrics) for facial analysis of static images. In order to speed image processing, the Emotrics source code was altered for batch processing by the senior author (JJG) using Sublime Text, a freely available Python code editor (**Supplemental Files**), based on open-access code for looping through directories (https://codegrepper.com/code-examples/python). The open-source Python algorithm utilizing ML libraries dlib and OpenCV libraries (hereafter called open-source algorithm) for dynamic facial detection in videos was available here (https://towardsdatascience.com/detecting-face-features-with-python-30385aee4a8e)^25^ and was adapted for batch processing for image and video analysis in a similar manner by the senior author (JJG) in Python v3.9.6 (**Supplemental Files**). Output of both files generated either a .jpg or a .mp4 file with overlaid 68 landmark dots (red-Emotrics, green-open-source Python algorithm (**Supplemental Files**) and a csv file containing x and y coordinates for each 68 landmarks per frame.

### Static Facial Analysis (Images)

Static images of 60 patients with normal facial function across the spectrum of FP severities (from mild to complete flaccid or synkinesis) were obtained from the MEEI database.^28^ Each patient had an image for the following 8 standard facial nerve exam movements: rest, eyebrow raise, gentle eye close, forceful eye closure, gentle smile, full effort smile, lip pucker, and lower lip depression. Open-source and the Emotrics algorithms were independently employed to detect key facial landmarks on all images (N=480). The resultant images of the open-source and Emotrics generated landmarks were arranged side by side in a Powerpoint slideshow. Each slide contained the landmark localization data from Emotrics and open-source for a single facial movement for one patient and had a pre-set 15 second timer. These images were separately analyzed by 3 independent blinded raters of varying clinical experience. The human eye was deemed to be the gold standard for landmark error detection. The presence of inconsistencies deviating from the reviewer’s perception of accurate landmark localization in five categories (facial contour, oral commissure, lips, eyelid, or eyebrows) was marked as a binary variable in an Excel spreadsheet.

### Dynamic Facial Analysis (Videos)

The open-source algorithm was used to generate the standard 68 landmarks on 30-50 second videos (frame rate of 30-60fps) of the same patient cohort performing the same 8 standard facial expressions from the MEEI database.^28^ The three independent graders watched each video together in real-time and independently noted the timing of significant landmark errors but were blinded to each other’s responses. An automated timer in the R studio program was utilized to obtain the start and stop points of any landmark inconsistencies to approximate the number of frames in which a landmark error was present. The time ranges spanning any landmark inconsistency in the same five facial categories were similarly noted in an Excel spreadsheet.

### Statistical analysis

To calculate error differences between the Emotrics and open-source algorithms for automated image analysis, the generated landmark coordinates 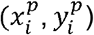 from the open-source Python algorithm was compared to the “ground truth” landmarks from Emotrics 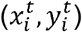 in order to compute the normalized root mean square error (NRMSE) and plot the Cumulative Error Distribution based on the methodology described by Johnston et al.^18^ The normalization factor was calculated based on the intercanthal distance (left eye 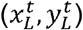 and right eye 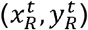, *d*_*norm*_, landmarks 39 and 42) to allow for comparison of different sized faces. NRMSE was converted to % for the cumulative error distribution and plotted in MATLAB (v R2021b, The Mathworks, Inc. 1994-2024). The Emotrics algorithm was considered the “ground truth” model as it represents the current standard for clinical facial landmark localization in the literature.^9,12,29,30^

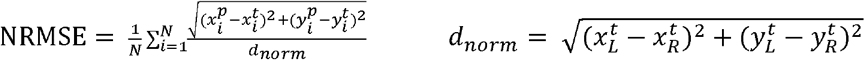

Additionally, separate error rates for Emotrics images and open-source algorithm images were calculated by facial feature type and FP severity based on individual graders’ responses. For each patient, we noted if there was at least one “major” error in each of the 8 standard facial expression frames and recorded it as a binary variable. These major errors were summed and divided by 8 to get an error rate, defined as major errors per frame, and these error rates per patient were averaged across the raters. These patient-specific error rates were then averaged based on FP type to generate average error rates for each FP type. These FP-specific rates were averaged again across all three reviewers. Similarly, average error rates were calculated for each facial feature. The averaged error rates of the Emotrics and open source algorithm-generated landmarks were compared using analysis of variance (ANOVA) tests.

For automated video analysis with the open-source algorithm, the duration of each recorded landmark inaccuracy was calculated based on the time ranges recorded by each independent grader and converted to number of frames by multiplying by the frame rate of each video. The number of frames with major errors were summed to approximate the total frames in which any landmark error was present in the video. The rate of landmark error was calculated as the number of frames with any landmark error divided by the total number of frames in the entire video (video length x frame rate). FP type and facial feature-specific error rates were similarly averaged across graders. These averaged error rates were compared between FP types between open-source generated images and videos with ANOVA tests. All statistical analysis was performed using the R statistical software (version 4.1.3).

## Results

### Static Facial Analysis (Images)

Cumulative error distribution of the NRMSE between the ground-truth Emotrics image analysis and the landmarks generated by the open-source ML-algorithm was most robust for participants with normal facial function and near-normal flaccid or mild synkinetic FP (**Figure 1**). The most significant discrepancies between Emotrics and the open-source ML-algorithm was found in cases of mild and moderate flaccid FP and near-normal and complete synkinetic FP (**Figure 1B-C**).

**Figure 1:**
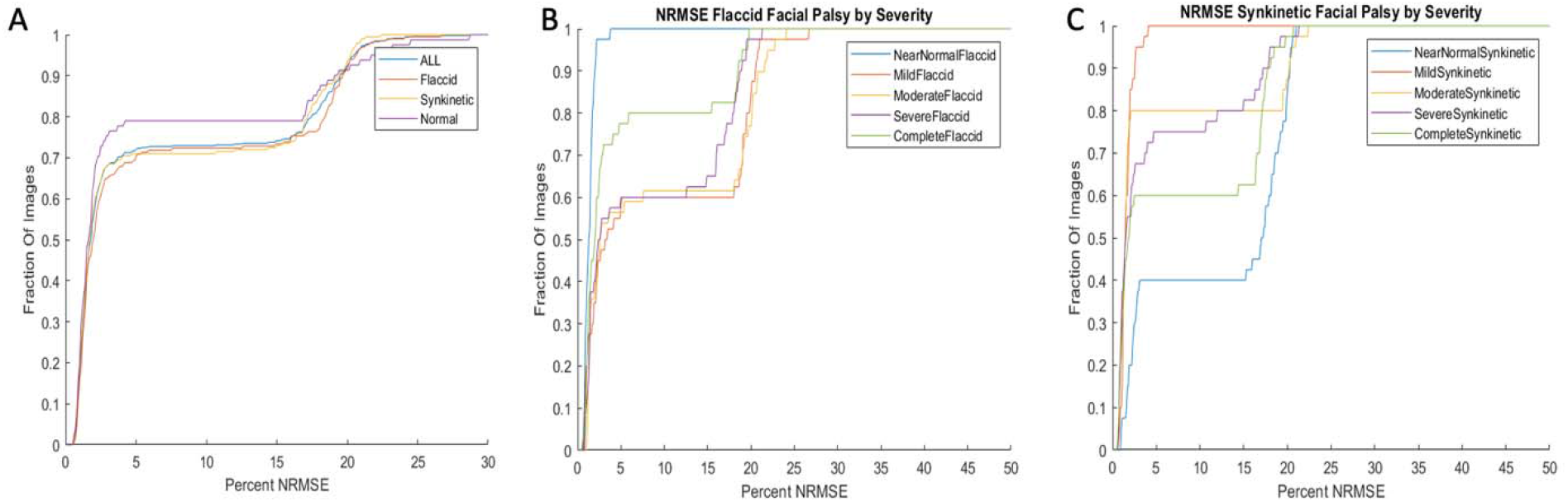
Cumulative error distribution of the NRMSE between the ground-truth Emotrics image analysis and the landmarks generated by the Python open-source ML-algorithm for A) all 8 standard facial movement images from the entire MEEI Facial Palsy database, separated by facial functional type, and by severity for B) flaccid facial palsy and C) nonflaccid or Synkinetic facial palsy.

An in-depth review of analyzed images revealed that both algorithms were prone to landmark inaccuracies. Examples of landmark errors by both Emotrics and the open-source algorithm are shown (**Figure 2**). In the clinician image error rate analysis, the open-source algorithm had slightly higher average error rates across all five facial features compared to Emotrics, but this difference was non-significant (p=0.76) (**Figure 3A**). Error rates [95% confidence intervals] followed similar patterns between the open-source algorithm and Emotrics; error rates were the highest for facial contours (open-source: 43.7% [41.2%, 44.9%] vs. Emotrics: 38% [36.1%, 39.6%]) and the lowest for eyebrows (open-source: 10.6% [8.9%, 12.8%] vs. 10.3% [8.5%, 12.5%]) in both algorithms. When analyzing error rates by FP severity, the open-source and Emotrics algorithms performed similarly (p=0.37) (**Figure 3B**). The highest average error rates for the open-source algorithm and Emotrics were in images of patients with severely flaccid FP (open-source: 88.3% [85.3%, 89.8%] vs. Emotrics: 83.3% [76.2%, 84.8%]). Notably, both programs had the same error average rate in severely synkinetic (open-source: 70.8% [68.6%, 71.4%] vs. Emotrics: 70% [69.1%, 72%]) and completely synkinetic patients (open-source: 70% [68.1%, 70.5%] vs. Emotrics: 69% [68.3%, 69.9%]). Although these differences were non-significant, the open-source algorithm had higher error rates than Emotrics across all other FP severities, except in near normal synkinetic patients (open-source: 80% [75.9%, 81.7%] vs. Emotrics: 82.5% [77.4%, 83.2%]) and mild synkinetic (open-source: 84% [83.8%, 85.7%] vs. Emotrics: 85.8% [84.3%, 86.8%]), where the open-source algorithm had a lower average error rate than Emotrics.

**Figure 2:** Sample of landmarks generated on static images by Emotrics and Python open-source machine learning algorithm. Both algorithms were prone to landmark inaccuracies. **\*Patient photos will be available in published manuscript**

**Figure 3:**
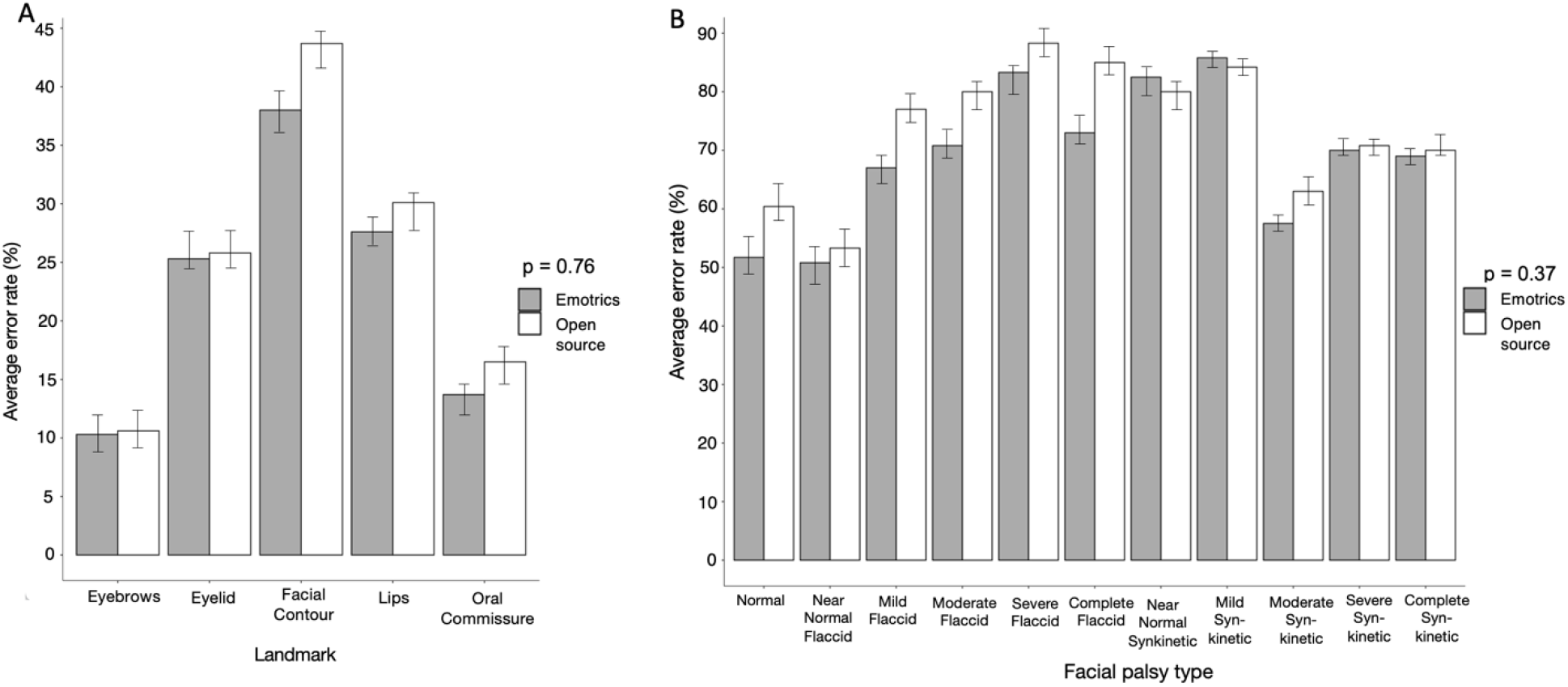
Comparison of Landmark Errors in Static Images of Facial Palsy by Emotrics and an open-source Python machine learning algorithm. (A) Examples of landmark errors from automated static image analysis by open-source Python machine learning algorithm (top right, inaccurate lips, oral commissure and facial contour) and by Emotrics (bottom, inaccurate lips, oral commissure and facial contour). Patient characteristics including facial hair and rhytids may play a role in landmark errors. There was no significant difference in average landmark error rates in static image analysis by Emotrics or the open-source Python algorithm when compared by (A) facial landmark and (B) facial palsy type and severity.

### Dynamic Facial Analysis (Videos)

We determined that it was feasible to employ the open-source algorithm^25^ for dynamic facial analysis in videos of FP patients (**Supplemental files**). Each 30-50 second video took 1-5 minutes to analyze on average on a Dell PC Windows 11 Pro featuring Intel(R) Core(TM) processor (i7-8850H CPU @ 2.60GHz 2.59 GHz) with 32GB of RAM. The average error rates were analyzed in the open-source algorithm-generated video landmarks across the entire MEEI database. The open-source algorithm performed most poorly on eyelid-tracking in videos, with eyelids having the highest average error rate of (20.7% [19.2%, 22.1%]) (**Figure 4A**). Lips had the next highest error rate (6.7% [5.4%, 6.7%]), followed by facial contour (1.5% [0.6%, 1.8%]), eyebrows (1.4% [0.8%, 1.9%]), and oral commissure tracking (0.34% [0.18%, 0.5%]). Based on FP type, the open-source algorithm had the highest error rates for mildly synkinetic patients (24.7% [22.3%, 25.1%]), followed by severely flaccid patients (19.7% [18.5%, 21.2%]) and completely flaccid patients (11.3% [10.4%, 11.8%]) (**Figure 4B**). The open-source algorithm had the lowest average error rates in nearly normal flaccid patients (7.1% [6%, 7.4%]) and mildly flaccid patients (8% [7.5%, 8.8%]). We did test the open-source algorithm on videos taken with a smartphone and found similar robust landmark accuracy as with the MEEI dataset videos (unreported data).

**Figure 4:**
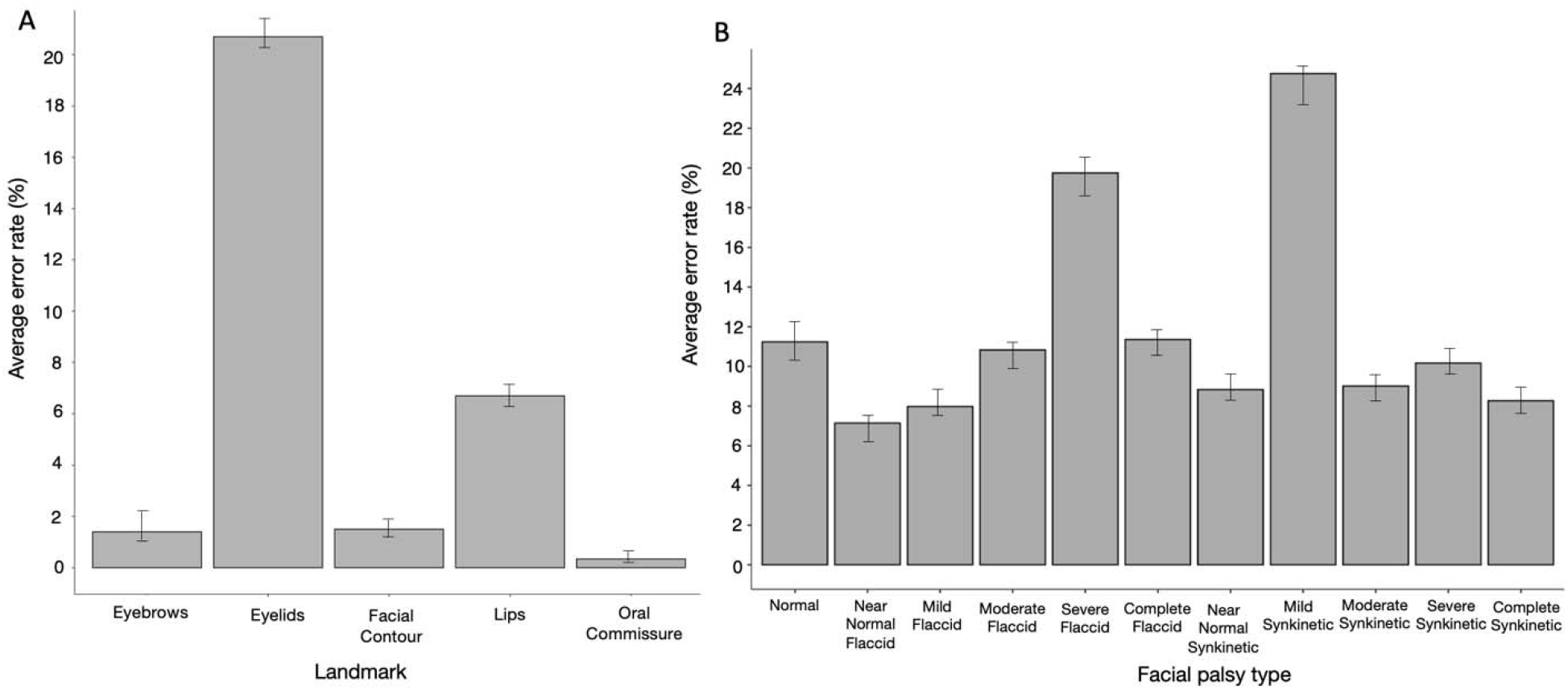
Average Landmark Error Rate of Dynamic Facial Analysis of Facial Palsy Patient Videos. Dynamic facial analysis of patient videos by the open-source Python algorithm revealed increased average error rates in certain facial landmarks such as the eyelids (A) and in particular facial palsy types such as severe flaccid and mild synkinetic patients (B).

### Landmark Error Rate in Static versus Dynamic Facial Analysis

When comparing error rates between image and video landmarks generated by the open-source algorithm of the MEEI database, videos had significantly lower average error rates than static images for the same patient, across all FP severity types (p < 0.001) (**Figure 5**). Completely flaccid patients had the largest reduction in average error rate and normal patients had the relatively lowest difference in error rate. Videos of facial nerve exams analyzed with the open-source algorithm yield an incredible increase in data compared to static images of facial nerve exams (images contain x and y coordinates for 68 landmarks for 8 frames = 1088 datapoints, videos contain x and y coordinates for 68 frames for a 30 second video (60fps) = 244,800 datapoints).

**Figure 5:** Dynamic Facial Analysis improves landmark accuracy. A significant reduction in the average landmark error rate was noted when the open-source Python algorithm was employed to analyze videos when compared to images of the standard facial nerve exam in patients with facial palsy across all types and severity levels (p<0.001). Sample of patient with images and a video of the standard facial nerve exam (**Supplementary Files**).

### Potential Applications of Dynamic Facial Analysis of Facial Palsy Videos

Future directions to quantify dynamic facial function from open-source ML-algorithm analysis of videos of FP are numerous. One example simply demonstrates plotting the time series of the y-coordinate of the oral commissure landmarks in a patient with normal facial function, complete flaccid and complete synkinetic FP (**Figure 6)**. Dynamic oral commissure synchrony in the y-axis is notable with a Pearson’s correlation coefficient of 0.99, but this linear trend varies with flaccid FP (Pearson’s r=0.75) and to a greater degree with synkinetic FP (Pearson’s r = 0.72). There are distinct patterns in facial movement that remain to be investigated with the aid of dynamic facial analysis with currently available ML-algorithms.

**Figure 6:**
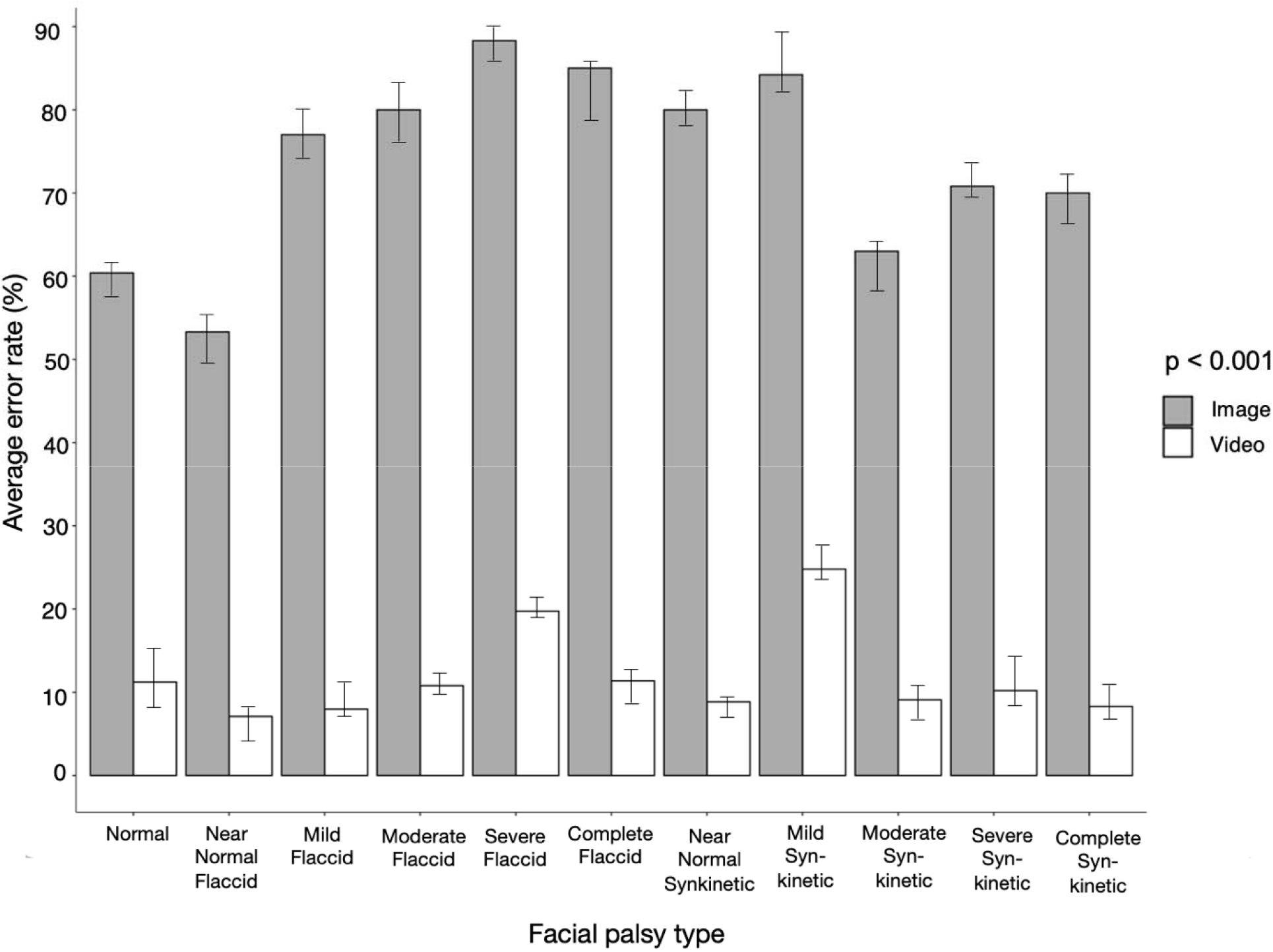
Potential Applications of Dynamic Facial Analysis of Facial Palsy Videos. We have found linear relationships in critical landmarks such as the oral commissure that become asynchronous and nonlinear in varying types of facial palsy. Dynamic oral commissure synchrony is displayed in the y-axis of patients with normal facial function, flaccid and synkinetic facial paralysis (top row, left to right). (Middle Row): Over the length of a video of a standard facial nerve exam, the right and left oral commissure y-coordinate is notably synchronous and symmetric in normal facial function (left) with a Pearson’s correlation coefficient of 0.99. This dramatically changes in patients with flaccid (middle) and synkinetic (right) facial paralysis. (Bottom Row): Scatter plots demonstrating the highly symmetric, linear relationship between the Right and Left oral commissure landmark in the y-axis in normal facial function(left) that drastically changes with facial palsy (middle, right). \***Patient photos will be available in published manuscript.**

## Discussion

Although our ability to read and interpret facial expressions is an evolutionary trait learned at a young age, our understanding of normal facial movements^31^ and ability to describe it mathematically is limited, particularly in the asymmetric and asynchronous movements of FP patients. However, should this challenge be met, we could significantly increase our understanding of facial nerve function and potentially decipher facial nerve regeneration in a manner not previously possible. A more precise evaluation of dynamic facial function would facilitate a better understanding of the spectrum of FP and improve guidance for and outcomes following facial reanimation surgery. With this study, we report several novel findings.

Firstly, the gold standard for facial analysis remains the human eye. We had used the Emotrics algorithm for static image analysis as the “ground truth” as it was re-trained on 1600 manually annotated images of patients with FP and represents the current standard for clinical facial landmark localization in the literature^9,12,29,30^. However, we found no significant differences in average error rate between both algorithms in static image analysis; both algorithms were susceptible to some landmark inaccuracies that the clinician raters picked up readily. Facial rhytids, facial hair, increased BMI, and patient movement are some of the suspected sources of error. Eyelid tracking was often lost in either modality particularly with forceful eyelid closure (likely due to the fact that the iris is a source of tracking in the dlib library). Patient effort is likewise something clinicians must note during documentation, particularly as FP patients may (consciously or not) minimize facial movements to mask their asymmetries. The solution to landmark errors in a traditional ML approach is to increase the size of the training dataset and to manually adjust landmarks; however, we agree with Guarin et al. that this approach is inefficient, prone to additional errors, and logistically difficult with the small datasets of FP images and videos. Future research into automated error detection, thresholds for video quality, and patient movement will further enhance the utility of ML-based analysis for dynamic facial function quantification. Consensus among providers who care for FP patients to determine the acceptable threshold error rate of ML-based dynamic facial analysis must be further investigated.

Secondly, contrary to contemporary beliefs regarding the limitations of ML algorithms for dynamic facial analysis of FP videos, we have found that not only was it feasible to use the latest open-source computer vision and ML algorithms available through Python (OpenCV, dlib) ^20,25,32^ on FP patients, but that landmark accuracy significantly improved compared to static image analysis. Because the volume of data available from a video of a standard facial nerve exam is orders of magnitude larger than static photography, we discovered a significant reduction in landmark error rate in videos compared to photos in a standardized data set across all FP severity types. Moreover, with the wealth of data video-based dynamic facial analysis can generate, patterns of facial movement have emerged from this analysis that have not previously been reported such as dynamic oral commissure synchrony. Video-based facial analysis facilitates visualization of dynamic facial movements and is the closest media format to in person assessment. Incorporation of dynamic facial analysis could provide an automated facial assessment to augment clinician evaluation, bolster quantitative outcomes tracking, and empower patients on the long road of facial nerve regeneration.

There are several limitations of this study that are important to recognize. The source dataset while high-quality and standardized, only contains 60 patients, and thus differences among subsets of FP severity may be over-emphasized. The standardized high-quality photo and video collection of the database may demonstrate reduced errors compared to patient-derived “in the wild” images and videos. Addressing landmark errors in such content represents a future path of research with the goal of facilitating patient-driven telemedicine facial nerve tracking of recovery and treatment outcomes. All facial image and video databases, including the MEEI Facial Palsy Photo and Video Standard Set, are limited in terms of patient age, racial and ethnic diversity. Future efforts to include racial and ethnic minorities in database participation is critical as FP can affect all groups and ages; however, the authors recognize that it is even more critical to protect the sensitive data for the patients who entrust us through multiple layers of data encryption, standard HIPAA privacy protections, and consent-taking, as data from FP patients is extremely sensitive.

## Conclusion

We report for the first time the feasibility and relative accuracy of a Python open-source machine learning algorithm for dynamic facial landmark tracking in FP videos and share the Python code. Due to the demonstrated superiority of landmark tracking using videos compared to images, we strongly encourage video documentation in addition to photo documentation of the facial nerve exam. We also report for the first time a linear relationship of the oral commissure with FP type that we term dynamic oral commissure synchrony. Further work into sources of error, dynamic facial patterns across the spectrum of FP, and developing a consensus among facial nerve clinicians regarding thresholds of error rate acceptable to clinical utility are needed to improve objective quantification of facial movement in FP.

---

**Supplemental Files:**
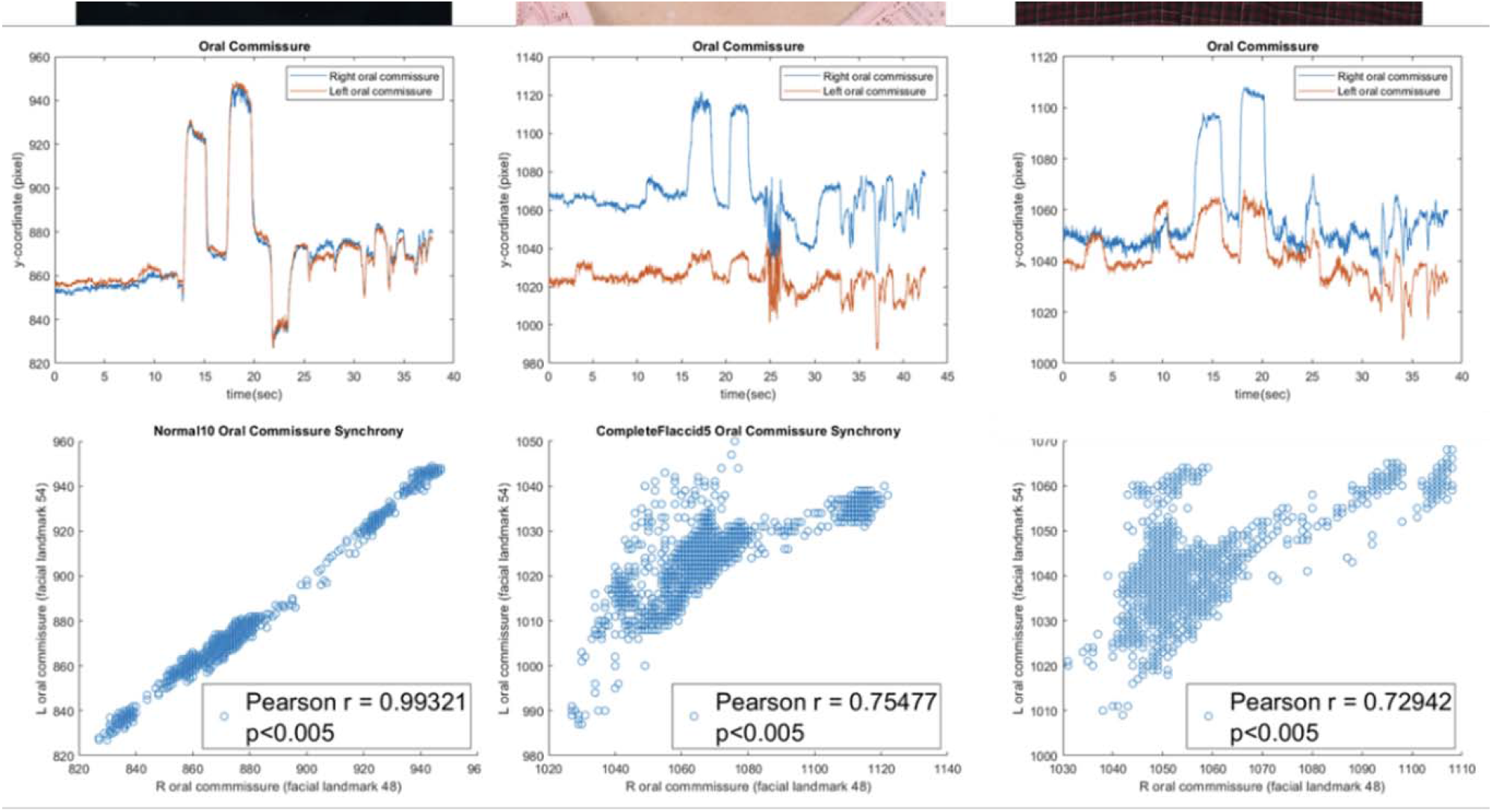
Files: not supplied in this version but will be available in the published manuscript. Python open source algorithm Python open source algorithm code.py for batch processing Landmarks generated by Python open source algorithm on a video of a normal function patient Coordinates of landmarks

## References

1. Hadlock T. Standard outcome measures in facial paralysis getting on the same page. JAMA Facial Plast Surg. 2016;18(2):85–86. doi:10.1001/jamafacial.2015.2095

2. Banks CA, Bhama PK, Park J, Hadlock CR, Hadlock TA. Clinician-graded electronic facial paralysis assessment: The eFACE. Plast Reconstr Surg. 2015;136(2):223e–230e. doi:10.1097/PRS.0000000000001447

3. Ng JH, Ngo RYS. The use of the facial clinimetric evaluation scale as a patient-based grading system in bell’s palsy. Laryngoscope. 2013;123(5):1256–1260. doi:10.1002/lary.23790

4. Gaudin RA, Robinson M, Banks CA, Baiungo J, Jowett N, Hadlock TA. Emerging vs time-tested methods of facial grading among patients with facial paralysis. JAMA Facial Plast Surg. 2016;18(4):251–257. doi:10.1001/jamafacial.2016.0025

5. Fattah AY, Gurusinghe ADR, Gavilan J, et al. Facial nerve grading instruments: Systematic review of the literature and suggestion for uniformity. Plast Reconstr Surg. 2015;135(2):569–579. doi:10.1097/PRS.0000000000000905

6. Revenaugh PC, Smith RM, Plitt MA, Ishii L, Boahene K, Byrne PJ. Use of objective metrics in dynamic facial reanimation a systematic review. JAMA Facial Plast Surg. 2018;20(6):501–508. doi:10.1001/jamafacial.2018.0398

7. Bylund N, Hultcrantz M, Jonsson L, Marsk E. Quality of Life in Bell’s Palsy: Correlation with Sunnybrook and House-Brackmann Over Time. Laryngoscope. 2021;131(2):E612–E618. doi:10.1002/lary.28751

8. Scheller C, Wienke A, Tatagiba M, et al. Interobserver variability of the House-Brackmann facial nerve grading system for the analysis of a randomized multi-center phase III trial. Acta Neurochir (Wien). 2017;159(4):733–738. doi:10.1007/s00701-017-3109-0

9. Guarin DL, Yunusova Y, Taati B, et al. Toward an Automatic System for Computer-Aided Assessment in Facial Palsy. 2020;22(1):42–49. doi:10.1089/fpsam.2019.29000.gua

10. Miller MQ, Hadlock TA, Fortier E, Guarin DL. The Auto-eFACE: Machine Learning-Enhanced Program Yields Automated Facial Palsy Assessment Tool. Plast Reconstr Surg. 2021;147(2):467–474. doi:10.1097/PRS.0000000000007572

11. Joseph Dusseldorp, Diego L. Guarin, Martinus M. Van Veen, Nate Jowett and Tah Hadlock TA. In the Eye of the Beholder: Changes in Perceived Emotion Expression after Smile Reanimation. Accept by Plast Reconstr Surg. 2018.

12. Dusseldorp JR, Van Veen MM, Guarin DL, Quatela O, Jowett N, Hadlock TA. Spontaneity Assessment in Dually Innervated Gracilis Smile Reanimation Surgery. JAMA Facial Plast Surg. 2019;21(6):551–557. doi:10.1001/jamafacial.2019.1090

13. Boonipat T, Asaad M, Lin J, Glass GE, Mardini S, Stotland M. Using artificial intelligence to measure facial expression following facial reanimation surgery. Plast Reconstr Surg. 2020:1147–1150. doi:10.1097/PRS.0000000000007251

14. Fuzi J, Meller C, Ch’ng S, Hadlock TM, Dusseldorp J. Voluntary and Spontaneous Smile Quantification in Facial Palsy Patients: Validation of a Novel Mobile Application. Facial Plast Surg Aesthetic Med. 2022;00(0). doi:10.1089/fpsam.2022.0104

15. Dusseldorp JR, Guarin DL, Van Veen MM, Miller M, Jowett N, Hadlock TA. Automated Spontaneity Assessment after Smile Reanimation: A Machine Learning Approach. Plast Reconstr Surg. 2022;149(6):1393–1402. doi:10.1097/PRS.0000000000009167

16. Hidaka T, Kurita M, Ogawa K, Tomioka Y, Okazaki M. Application of Artificial Intelligence for Real-Time Facial Asymmetry Analysis. Plast Reconstr Surg. 2020;146(2):243e–245e. doi:10.1097/PRS.0000000000007035

17. Guarin DL, Bandini A, Dempster A, et al. The Effect of Improving Facial Alignment Accuracy on the Video-based Detection of Neurological Diseases. J Biomed Heal Informatics. 2017;XX(Xx):1–9. doi:10.36227/techrxiv.12950279.v1

18. Johnston B, Chazal P de. A review of image-based automatic facial landmark identification techniques. Eurasip J Image Video Process. 2018;2018(1). doi:10.1186/s13640-018-0324-4

19. Sagonas C, Tzimiropoulos G, Zafeiriou S, Pantic M. 300 faces in-the-wild challenge: The first facial landmark Localization Challenge. Proc IEEE Int Conf Comput Vis. 2013:397–403. doi:10.1109/ICCVW.2013.59

20. King DE. Dlib-ml: A machine learning toolkit. J Mach Learn Res. 2009;10:1755–1758.

21. Diego L Guarin, Joseph Dusseldorp, Tessa Hadlock NJ. A Machine Learning Approach for Automated Facial Measurements in Facial Palsy An ongoing problem in the management of facial neuromo-. JAMA Facial Plast Surg. 2019;20(4):2018–2020. doi:10.1007/s002669900252

22. Thies J, Zollhöfer M, Stamminger M, Theobalt C, Nießner M. Face2Face: Real-time face capture and reenactment of RGB videos. Commun ACM. 2019;62(1):96–104. doi:10.1145/3292039

23. Guarin DL, Yunusova Y, Taati B, et al. Toward an Automatic System for Computer-Aided Assessment in Facial Palsy. Facial Plast Surg aesthetic Med. 2020;22(1):42–49. doi:10.1089/fpsam.2019.29000.gua

24. Guarin, Diego L; Dusseldorp, Joseph R; Hadlock, Tessa A; Jowett N. A Machine Learning Approach for Automated Facial Measurements in Facial Palsy. JAMA Facial Plast Surg. 2018;20(4):335–337. doi:10.1007/s002669900252

25. Martinez JC. Detecting Face Features with Python. Live Code Stream. https://livecodestream.dev/post/detecting-face-features-with-python/.

26. Shen J, Zafeiriou S, Chrysos GG, Kossaifi J, Tzimiropoulos G, Pantic M. The First Facial Landmark Tracking in-The-Wild Challenge: Benchmark and Results. Proc IEEE Int Conf Comput Vis. 2016;2015-February:1003–1011. doi:10.1109/ICCVW.2015.132

27. Malm, IJ, Albathi, M, Byrne, P, Ishii, M, Ishii, L, Boahene K. Facial Asymmetry Index: Validation and Applications in Various Smile Restoration Techniques. Facial Plast Surg. 2018;34(4):381–383.

28. Greene JJ, Guarin DL, Tavares J, et al. The spectrum of facial palsy: The MEEI facial palsy photo and video standard set. Laryngoscope. 2020;130(1):32–37. doi:10.1002/lary.27986

29. Greene JJ, Tavares J, Guarin DL, Hadlock T. Clinician and Automated Assessments of Facial Function Following Eyelid Weight Placement. JAMA Facial Plast Surg. 2019;02114:1–6. doi:10.1001/jamafacial.2019.0086

30. Perez PB, Gunter AE, Moody MP, et al. Investigating Long-Term Brow Stabilization by Endotine-Assisted Endoscopic Brow Lift with Concomitant Upper Lid Blepharoplasty. Ann Otol Rhinol Laryngol. 2021;130(10):1139–1147. doi:10.1177/0003489421997653

31. Ekman P, Oster H. Facial Expressions +316 of Emotion. Annu Rev Psychol. 1979;30:527–554.

32. Meijerink C. Facial Landmark Detection Under Challenging Conditions. 2021. http://essay.utwente.nl/86867/.

